# Understanding the genetic diversity of bacteria isolated from across the Atacama Desert

**DOI:** 10.1101/2025.08.25.672134

**Authors:** Nicole Cavanaugh, Alicyn Reverdy Pearson, Elliot Ingraham, Elizabeth Amorelli, Austen Herlihy, Nathan Thewedros, Matteo Couto Frignani, Marcello Twahirwa, Carlos Riquelme, Yunrong Chai, Veronica Godoy-Carter

## Abstract

Despite being one of the driest and harshest deserts on Earth, the Atacama Desert is home to a variety of bacterial life. Microorganisms that reside here may have developed adaptations to help them survive this unique environment. In this study, we used bioinformatic and genetic methods to assess the abundance of phyla that are present in this environment and what types of adaptations individual bacteria have obtained. To assess bacterial diversity, we used 16S rRNA sequencing on soil samples and determined the relative composition of different phyla and archaea at sixteen locations. A selection of eight cultivatable organisms which produce pigments were subjected to whole genome sequencing (WGS). Using these sequences, we screened for stress-tolerance capabilities including pigment production pathways, biofilm-related genes, antibiotic production, and genome stability. We found that all strains we sequenced are predicted to produce bioactive compounds. We also found that the pigments that these bacteria produce have antioxidant, iron and ion chelating, and/or antibiotic properties. This characterization allows us to assess adaptive strategies of bacteria which is important in the fields of agriculture, biotechnology and health.

## 1. Introduction

Even the harshest landscapes can be home to diverse microbial life. Microbes have been identified as far as humans have been able to sample, from the deep depths of the sea floor to the Arctic to near the centers of active volcanoes (1–3). Understanding how bacteria survive stressful conditions is essential to many areas including agriculture, biotechnology, and human health.

Organisms that are capable of living in harsh conditions are called extremophiles. Extremophiles are known to develop adaptations to help them survive and thrive. Examples include structural changes in the DNA such as G-quadruplex formation (G4), and/or high GC content, pigment production, biofilm formation, production of unique secondary metabolites, and antibiotic resistance mechanisms, among others. Some microorganisms produce pigments that protect them from damage by UV rays, such as melanin or carotenoids (4,5). Many bacteria are capable of forming biofilms, which are multicellular communities of bacteria bound together by a self-produced matrix (6). The matrix provides many benefits to the community including providing a medium for concentrating and sharing nutrients, adding a protective barrier to protect the community from antimicrobials, maintaining hydration, and protecting the cells from UV radiation (6–9). It is also beneficial for bacteria to produce compounds that inhibit growth of other organisms to protect their community and resources (10). On the other hand, many organisms also develop resistance mechanisms to help them withstand inhibitory compounds produced by other organisms (11,12).

In addition to producing unique molecules to aid in survival, characteristics of an organism’s DNA itself can be an adaptation. DNA is made up of nucleotide pairings, where adenine (A) pairs with thiamine (T) and guanine (G) pairs with cytosine (C) (13). DNA that is higher in GC content may have stronger DNA stability due to more favorable nucleotide stacking patterns (14). It has also been shown in plants that individuals that live in cold or dry climates are more likely to have a high GC content (15). Sections of DNA that contain several runs of guanines are sometimes capable of forming G-quadruplexes (G4’s), which is a non-helical secondary structure of DNA (16). G4’s form in a wide variety of organisms, including bacteria, and serve a variety of functions. While investigating bacterial G4 functions is a newly popular field, it had been shown that G4’s in bacteria often concentrate near promoter regions (17). In particular, RecA-dependent DNA repair pathways in *Deinococcus radiodurans* and *Deinococcus geothermalis*, known for being able to withstand lethal doses of radiation, are partially regulated by G4’s (16).

One environment that is a high priority for scientists studying extremophiles is the Atacama Desert, located in northern Chile, South America. The Atacama Desert is recognized as one of the oldest, driest, and most UV-irradiated deserts on Earth (18,19). The desert spans approximately 128,000km^2^ and sits at 2,000-5,000 meters above sea level, limiting available oxygen and causing high levels of UV radiation (20,21). Additionally, the desert lies between two mountain ranges, the Andes Mountains and the Chilean Coast mountain range, which severely limit precipitation in the area (18,19). For these reasons, the Atacama Desert continues to be an important site for studying how microorganisms can adapt to harsh environments. Over the course of several years, we have surveyed and sampled a variety of locations across the desert with the goal of understanding how bacteria live there and what adaptations they have specifically developed to survive this unique environment.

In July 2018 we performed global 16s sequencing on soil samples across 18 locations with the goal of understanding what species lived across the Atacama Desert. To enhance this work, in 2022 and 2024, we isolated and whole genome sequenced bacteria from a variety of locations throughout the desert. We selected strains that produce pigments in different conditions. Since pigment production can often be seen visually, pigment formation is a great starting point for looking at sequencing candidates for well-adapted bacteria. Organisms may produce pigments for a variety of reasons including ultra-violet (UV) protection, antioxidant purposes, biofilm control, and sequestering molecules, to name a few. Using our genomic sequences, in addition to identifying the pigments the isolates produce, we can also identify other mechanisms that are beneficial for survival in inhospitable environments, including production of antimicrobial and bioactive compounds, key biofilm formation genes, and cell communication genes.

The knowledge gained from this genetic survey will enable evolutionary understanding of key survival strategies in bacteria. Insight into how bacteria handle stress is relevant in many fields including agriculture, biotechnology and health.

## 2. Materials and Methods

### 2.1. Metagenomic Methods

#### 2.1.1. Sample Collection, Preparation, and Storage

Environmental samples were collected during the months of May and June of 2018. (Figure 1) At each location, soils were collected from between 1-5 cm depth using sterile tools. Samples were stored in sterile 2mL microcentrifuge tubes. The date and exact coordinates were recorded at each sampling location (Table S1). Samples were kept at room temperature while in the field. Once brought to the laboratory, 20% (v/v) glycerol was added to the samples. Samples were stored at −80º C until thawed to prepare for 16SrRNA metagenomic analysis.

**Figure 1.**
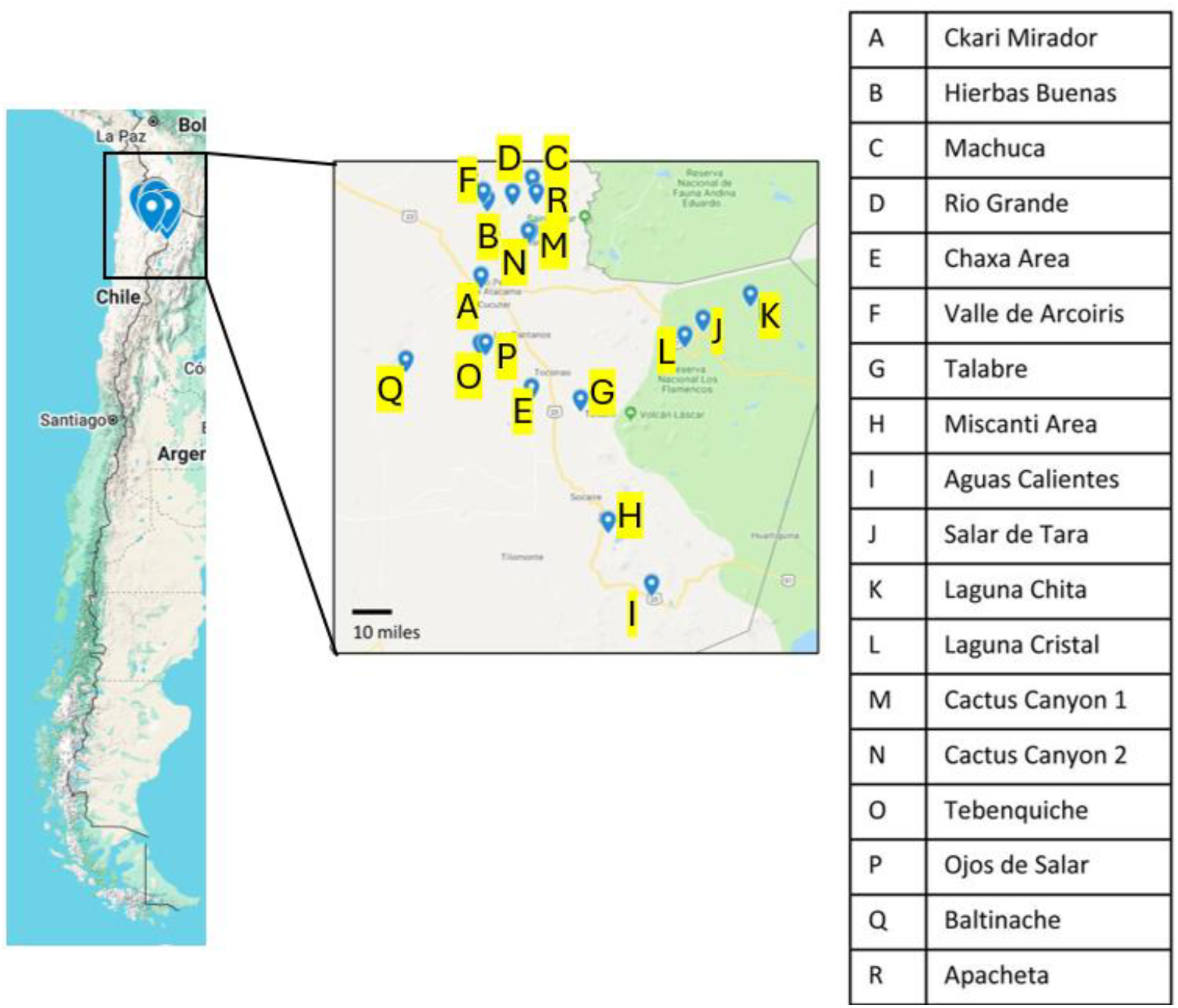
2018 Soil Sampling Map. On a map of Chile (left) locations where samples were collected in 2018 for metagenomic analysis are highlighted. Sampling locations are listed (right) and noted on a zoomed-in section of the map showing part of the Atacama Desert (center).

#### 2.1.2. 16SrRNA Metagenomic Analysis

Approximately 1g of soil sample was added to UBiome microbiome sequencing kit tubes (UBiome, San Francisco, CA). Samples were processed and analyzed with the ZymoBIOMICS® Service: Targeted Metagenomic Sequencing (Zymo Research, Irvine, CA).

#### 2.1.3. Targeted Library Preparation

The ZymoBIOMICS ® DNA Miniprep Kit (Zymo Research, Irvine, CA) was used for DNA extraction. Bacterial 16S ribosomal RNA gene targeted sequencing was performed using the Quick-16S™ NGS Library Prep Kit (Zymo Research, Irvine, CA). The bacterial 16S primers amplified the V3-V4 region of the 16S rRNA gene using primers custom-designed by Zymo Research to provide the best coverage of the 16S gene while maintaining high sensitivity. The sequencing library was prepared using a Zymo-developed library preparation process in which PCR reactions were performed in real-time PCR machines to control cycles and limited PCR chimera formation. The final PCR products were quantified with qPCR fluorescence readings and pooled together based on equal molarity. The final pooled library was cleaned up with the Select-a-Size DNA Clean & Concentrator™ (Zymo Research, Irvine, CA), then quantified with TapeStation® and Qubit®.

#### 2.1.4. Metagenomic 16s Sequencing

The final library was sequenced on Illumina ® MiSeq™ with a v3 reagent kit (600 cycles). The sequencing was performed with >10% PhiX spike-in. Amplification was compared to a negative control standard.

#### 2.1.5. Bioinformatic Analysis

Unique amplicon sequences were inferred from raw reads using the Dada2 pipeline (22). Chimeric sequences were also removed with the Dada2 pipeline. Taxonomy was assigned using Uclust from Qiime v.1.9.1 with Greengenes 16S database as reference. Visualization and Analysis of Microbial Population Structure (VAMPS) program was used for all subsequent diversity, abundance, and cluster plot analyses (23). Alpha diversity, or richness, was quantified as the number of unique sequences found in the sample. Phyla and genera relative abundance were calculated as the percent number of sequences out of the total number of sequences in the sample. The cluster plots were calculated based on Morisita Horn parameters at the genus level.

### 2.2. Whole-Genome Methods

#### 2.2.1. Sampling and Bacterial Isolation

To study how specific bacteria can handle stress, we started by isolating a variety of bacterial colonies from the desert soil. Samples were collected in July 2022 and July 2024. At each sampling location, soils and other environmental samples were collected from between 1-5 cm depth using sterile tools. (Figure 2) Samples were stored in sterile 2mL microcentrifuge tubes. The date and exact coordinates were recorded at each sampling location (Table S2). Samples were kept at room temperature while in the field. Once brought to the laboratory, bacterial colonies were isolated after plating 100 µL of a water/sample slurry on Reasoner’s 2A (R2A) agar medium and incubating plates at 28 ºC for 48-72 hours. This isolation yielded bacteria of many morphologies and colors. Each isolate was stocked in 20% (v/v) glycerol in water and stored at −80º C.

**Figure 2.**
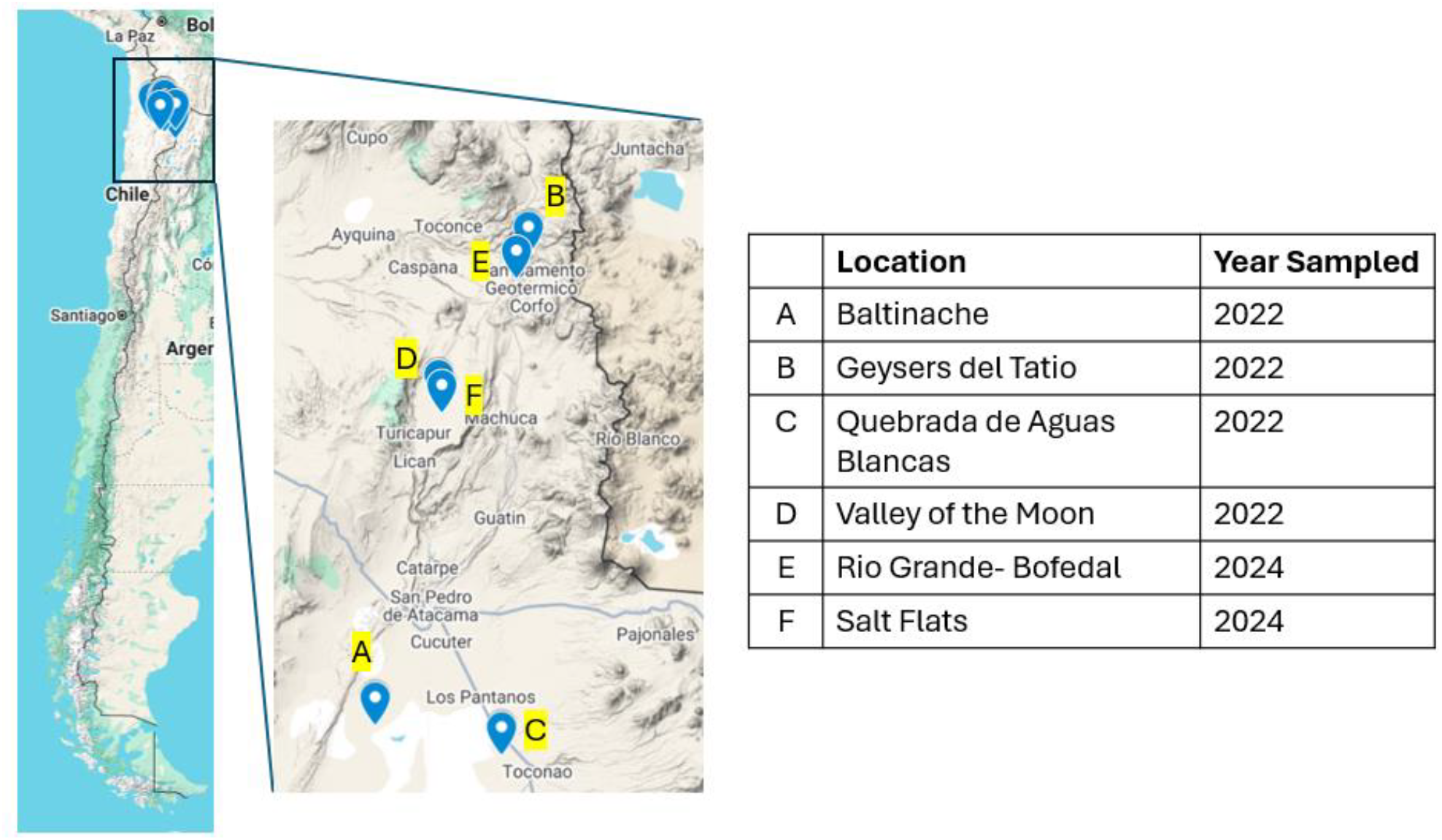
Sampling locations 2022 and 2024. On a map of Chile (left) sampling locations from 2022 and 2024 are labeled. In 2022 samples were collected from Baltinache (A), Geysers del Tatio (B), Quebrada de Aguas Blancas (C), and Valley of the Moon (D). In 2024 samples were collected from Rio Grande-Bofedal (E) and from salt flats near the Valley of the Moon (F).

#### 2.2.2. Biofilm Growth

To test biofilm formation in isolates that were phenotypically like *Bacillus subtilis*, strains were grown for 4 hours at 37 ºC to OD600 = 1.0 in LB broth in shaking conditions. 2 µl of culture was spotted on LBGM agar plates- biofilm-inducing plates where LB is supplemented with 1% glycerol and 0.1 mM MnSO4 (24). Colony biofilms were grown at 30 ºC for 72 hours before imaging.

A selection of isolates that produce pigments in a variety of conditions became candidates for whole genome sequencing.

#### 2.2.3. Genome Sequencing and Assembly

The genomes of three isolates, 0102A, 0209A, and 0909A, were sequenced by Illumina MiSeq (Illumina) as described in Cavanaugh, et al., 2024. Details about sequencing and assembly can be found in the publication (25).

For isolates 1020B, 0516A, 0819A, Iso1_2024, and Iso2_2024, genomic DNA was isolated from 2-mL R2B cultures using Promega’s Wizard Genomic DNA Purification Kit (Promega, USA), following the provided instructions for Gram-positive bacteria. Purified DNA was quantified using a NanoDrop spectrophotometer (Thermo-Fisher).

Purified gDNA was sequenced by Oxford Nanopore long-read sequencing (Oxford Nanopore Technologies) by Plasmidsaurus. Libraries were prepared using the V14 Ligation Sequencing Kit and sequencing was performed using the MinION R10.4.1 flow cell (Oxford Nanopore Technologies). Before assembling the genomes, the 5% lowest quality reads were removed by Fitlong v0.2.1. (26). Reads were downsampled to 250 Mb by Fitlong and a rough assembly was generated with Miniasm v0.3. (27). Reads were further downsampled to 100x coverage when coverage was over 100x. Quality reads were assembled using Flye v2.9.1 (28) and cleaned using Medaka v1.8.0 (Oxford Nanopore Technologies).

#### 2.2.4. Construction of Phylogenetic Tree Using autoMLST

A phylogenetic tree was generated using the autoMLST2 web server using denovo mode (29). autoMLST2 was run using default parameters.

#### 2.2.5. antiSMASH Analysis

Biosynthetic Gene Cluster (BGC) analysis was performed on all genomes using antiSMASH v8.0.2 using a relaxed detection strictness (30). FASTA files were used and KnownClusterBlast, TFBS analysis, activesitefinder, SubClusterBlast, and RREFinder analyses were selected from the “extra features” menu. Secondary metabolite-producing regions found in each genome were recorded.

#### 2.2.6. Bioinformatic Analysis of *Kocuria sp*. Strains’ Carratenoid BGCs

For individual identification of carotenoid synthesis genes, DIAMOND v2.0.15 alignment software was utilized to align translated protein sequences against the RefSeq Bacterial Protein Database (REFSEQBP) (31). Additionally, 0819A’s genome was aligned against all known *Kocuria rhizophila* proteins (KRAP), Iso2_2024’s genome was aligned against all known *Kocuria turfanensis* proteins (KTAP), and 1020B’s genome was aligned against all known *Kocuria oceani* proteins (KOAP)^6^. For REFSEQBP alignments, alignments were performed using very relaxed parameters, compared to the more stringent parameters used for the KTAP, KRAP, and KOAP alignments. For each strain, the top 15,000 best aligned proteins were selected and sorted for carotenoid biosynthesis genes. A pipeline was created using Bash and Python languages in the Linux environment, creating scripts to download databases, converting them to DIAMOND format, performing the DIAMOND alignments, assigning protein names to taxonomic IDs, arraying the data into an interpretable format, and back calculating genome positions of hits.

#### 2.2.6. Analysis by G4 Hunter

Genome GC content and g4 quadruplex (G4) content was assessed by the G4 Hunter web application, available through DNA Analyser (32). Genome FASTA files were uploaded to G4 Hunter and analyzed using a window size of 25 nt and a threshold of 1.2. The estimated number of G4’s, G4 frequency per thousand base pairs (kbp), GC content, and genome size were recorded.

## 3. Results

### 3.1. 16s Metagenomics

To assess the microbial community composition of the samples, Illumina 16S rRNA sequencing was used directly on soil samples from the 18 sampling locations and analyzed for its Archaea and Bacteria composition. Importantly, the soil samples from Ojos de Salar and Baltinache did not provide reliable data when compared to the negative control and could not be analyzed any further. This indicates that the soil composition was such that it did not permit the identification of microbial life with the test used. Community composition was successful in the remaining 16 soil samples.

The focus of our search was on bacterial diversity. Looking at the number of different taxa in a sample, the alpha diversity, a measurement of the richness, ranged from 184 to 652 taxa (Figure 3A). Overall, this is low compared to other environments. Sea water has an estimated 10,000 species richness, the temperate forest and grasslands soil has a few thousand species, and the gut microbiome has about one thousand species(33–36). As predicted, our results suggest that the extreme environment has a significantly lower number of unique taxa detected using the direct sequencing method. Unfortunately, it is difficult to make direct comparisons with other studies as the sequencing methods have changed dramatically since these started.

**Figure 3.**
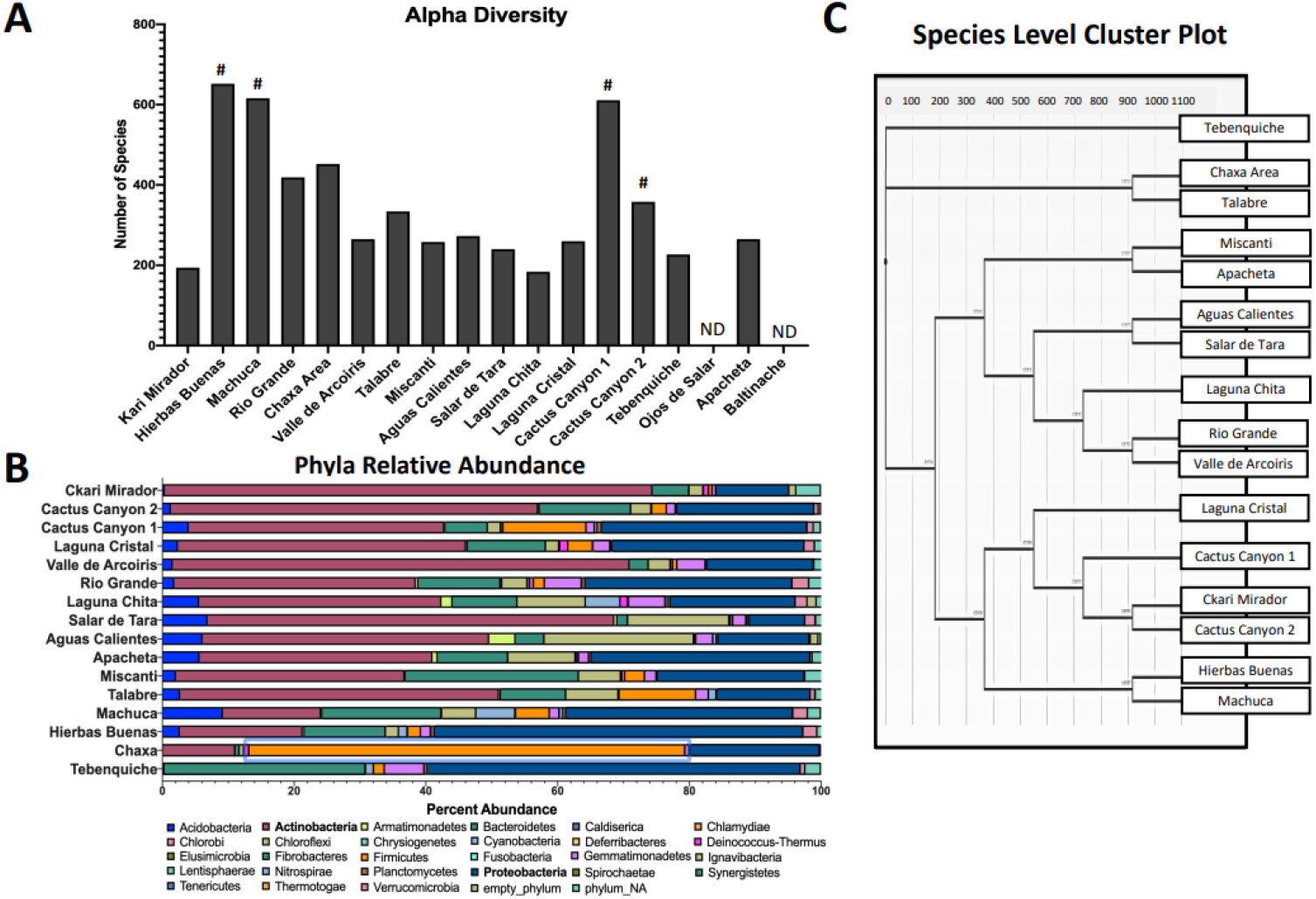
Illumina 16S rRNA sequencing provides a snapshot of bacterial diversity. (A) Quantification of alpha diversity soil samples. Each bar represents the number of unique taxa detected in each soil sample. Pound signs indicate locations of higher plant life. There was no detection of sequences in Ojos de Salar or Baltinache (ND). (B) Bar chart of the phyla relative abundance within each sample. Each color represents a different phylum, and the length of the bar represents the percent abundance within that sample. Actinobacteria (pink) and Proteobacteria (dark blue) were the most abundant phyla. (C) Cluster plot dendrogram grouping samples based on genus similarities under Marisita-Horn analysis. Length of bar represents degree of similarity. Tebenquiche was the most different from the rest of the samples.

We were curious to find out what was the percentage of Archaea in our samples as many are well known extremophiles (37,38). We found that 1.1% of the community was Archaea (Figure 4). This is comparable to levels found in the other hyper-arid desert soils (37,39).

**Figure 4.**
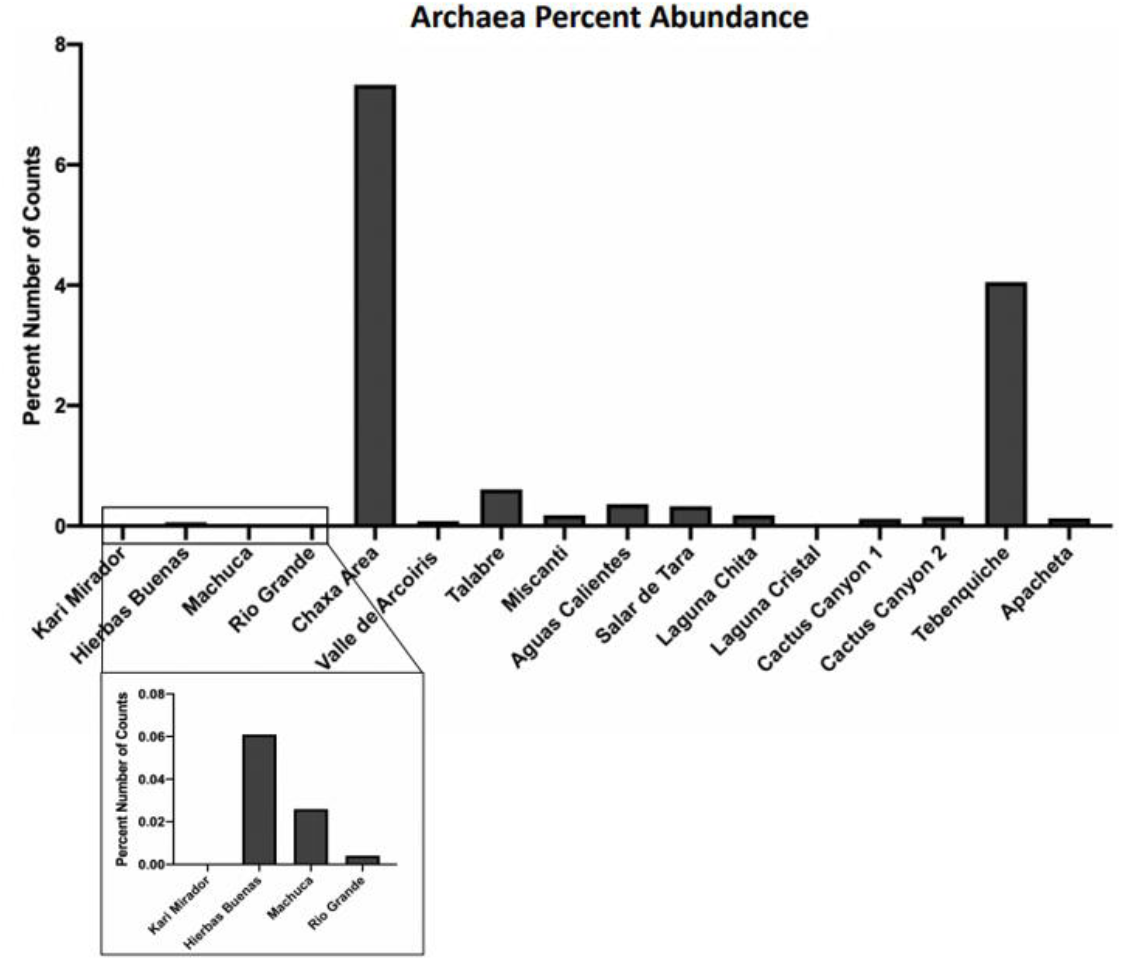
Archaea make up 1.14% of the total community diversity. The percent number of Archaea sequences of total samples was calculated from the Illumina 16S rRNA sequencing data set for each location. The total makeup of all detected sequences was 1.14% Archaea. Chaxa and Tebenquiche had the highest abundance of Archaea. Ckari Mirador and Laguna Cristal contained no detection of Archaea (box is a zoomed in look at the values).

We find that locations that have more plant life, had a greater richness (Figure 3A). Hierbas Buenas (Good Herbs) has a comparatively large abundance of low grass and bushes, and accordingly, had the largest richness at 652 taxa (Figure 3A). Conversely, the locations with the driest and most barren soil had the lowest richness as seen at Laguna Chita, Ckari Mirador, Tebenquiche, Salar de Tara, and Valle de Arcoiris with 184, 194, 227, 240, and 265 taxa, respectively (Figure 3A). As expected, some of these locations also had the highest levels of Archaea (Figure 4) which are well known extremophiles (37,38). The correlation of “plant-rich” areas containing higher bacterial richness is expected due to the strong symbiosis and known importance that the bacterial rhizobiome has for plant growth and survival.

The two dominant phyla across all locations were Actinobacteria at 35.9% (pink bars) and Proteobacteria (dark blue bars) at 26.7% (Figure 3B). Our results are consistent with other studies that have investigated the bacterial content in the Atacama Desert (39–41). Other studies have also found Bacteroidetes and Cyanobacteria to be the dominant phyla, however they were sampling water, while we sampled soil (42–44). Bacteroidetes is the third most abundant phylum across all the samples at 11.3% abundance (Figure 3B).

To compare the bacterial diversity across the different sampling locations, we generated a genus level cluster plot (Figure 3C). The salt flat, Tebenquiche, was the outlier group and the other salt flat, Chaxa, was in another outlier grouping with Talabre. The clustering of Apacheta and Miscanti is most likely due to their high altitude, shrub-containing characteristics. Salar de Tara and Aguas Calientes are closely located and near high altitude lakes (Figure 1). Rio Grande and Valle de Arcoiris are two very dry and barren locations. The clustering based on location similarities highlights that the genera are most likely selected by the environmental conditions.

### 3.2. Genomic Evidence

After bacteria were isolated from environmental samples on R2A medium, several appeared to produce pigments. Isolates 1020B, 0819A, and Isolate 2 displayed strong yellow and orange pigmentation after growing on R2A medium for 3 days. (Figure 5C-E). Two isolates also appeared to secrete blue and green pigments. (Figure 5A-B).

**Figure 5.**
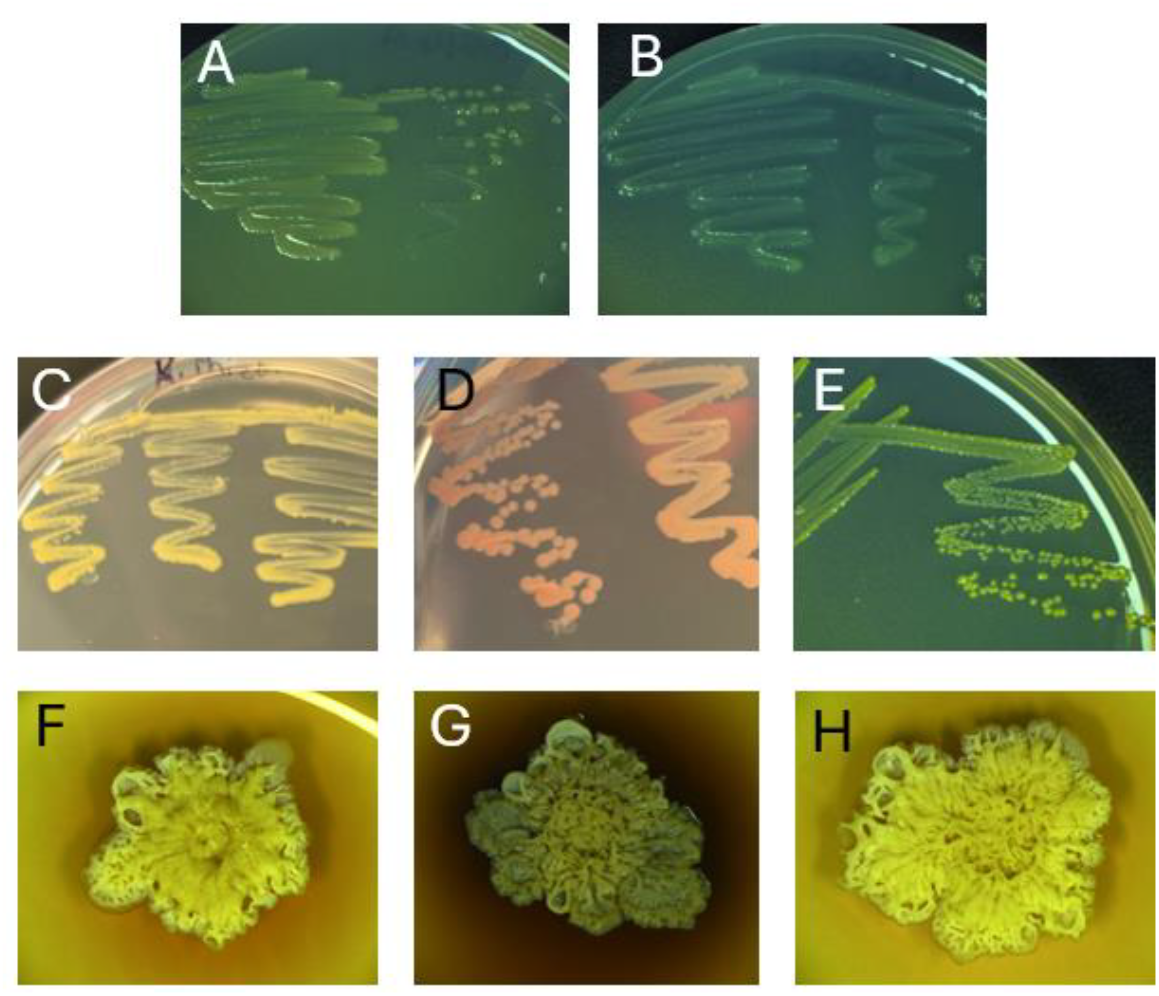
Selected isolates for WGS produce pigments. 0516A secretes a faint green pigment (A), Iso2_2024 secretes a blue pigment (B), 0819A and 1020B produce yellow pigments (C and E), and Iso2_2024 produces an orange pigment (E) on R2A medium. 0102A, 0209A, and 0909A secrete red-brown pigments when grown on LBGM biofilm-inducing medium (F-H).

Three isolates appeared to have similar colony morphology to *Bacillus subtilis-* 0102A, 0209A, and 0909A. It is known that *B. subtilis* produces a pigment called pulcherrimin late in the biofilm formation cycle, regulating biofilm aging and disassembly (45). When these isolates were grown on LBGM biofilm inducing medium, they produced a red-brown pigment that resembled pulcherrimin. (Figure 5F-H) These eight isolates were selected to be subjected to whole genome sequencing (WGS). (Figure 5)

After sequencing, all resulting genome FASTA files were aligned using autoMLST2 (29). autoMLST2 uses multiple loci across the genome, focusing on housekeeping genes to provide higher resolution, to identify the most closely related species for each sequence by generating a multi-locus species tree. (Figure 6) autoMLST2 automatically identified related genome sequences to create the species tree de novo. autoMLST2 was able to identify the identities of most isolates to the species level. Iso1_2024 was identified as *Pseudomonas aeruginosa*, 0516A was identified as *Pseudomonas rhodesiae*, 0819A was identified as *Kocuria rhizophila*, 1020B was identified as *Kocuria oceani*, Iso2_2024 was identified as *Kocuria turfanensis*, and 0909A was identified as *Bacillus mojavensis*. (Table 1) All identifications were made with a high level of confidence, according to the Mash distance (46), estimated average nucleotide identity (ANI), and P-value. The top hits for 0102A and 0209A identification were “*Bacillus subtilis* group”, rather than being matched directly with a species. The top three most similar species matches are listed in Table 2. Both strains are most closely related to *Bacillus inaquosorum*. (Table 2)

**Table 1.**
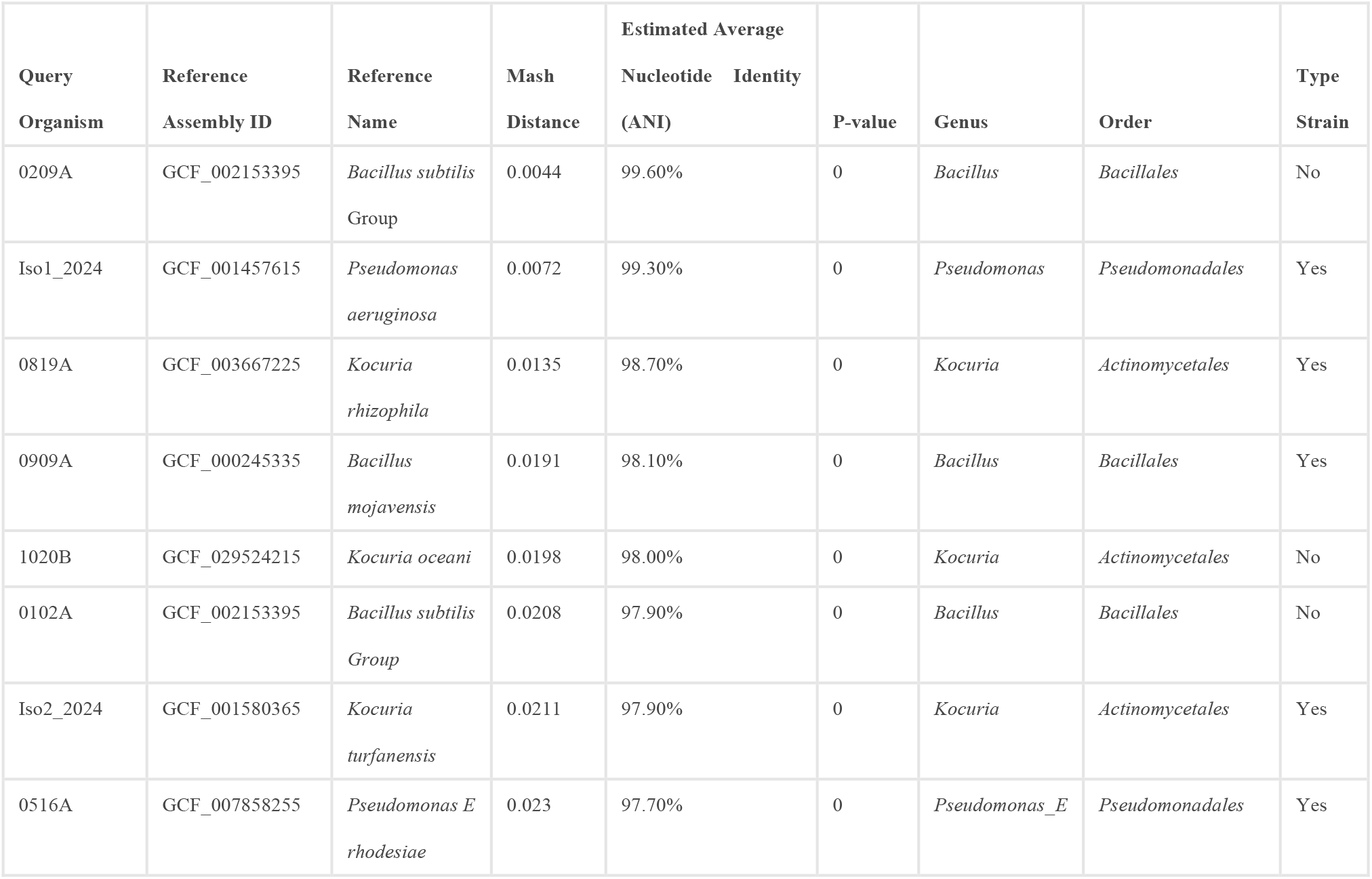
Species Identification for isolates with sequenced genomes by autoMLST2. Query organisms, those sequenced in this study, were aligned by autoMLST2 against a large database of sequences to find the most closely related species for each. Each organism’s top match is listed in the “Reference Name” column and the sequence ID is listed under “Reference Assembly ID”. The genus, order, and whether or not the match was a type strain, are listed in columns at the right. The quality of the match is assessed by Mash distance (a similarity assessment utilizing mutation distance and P-value significance), average estimated nucleotide identity (ANI-percent nucleotide similarity between two sequences), and P-value (statistical chance that the match was made by chance).

| Query<br>Organism | Reference<br>Assembly ID | Reference<br>Name | Mash<br>Distance | Estimated Average<br>Nucleotide Identity<br>(ANI) | P-value | Genus | Order | Type<br>Strain |
| --- | --- | --- | --- | --- | --- | --- | --- | --- |
| 0209A | GCF_002153395 | *Bacillus subtilis*<br>Group | 0.0044 | 99.60% | 0 | *Bacillus* | *Bacillales* | No |
| Iso1_2024 | GCF_001457615 | *Pseudomonas aeruginosa* | 0.0072 | 99.30% | 0 | *Pseudomonas* | *Pseudomonadales* | Yes |
| 0819A | GCF_003667225 | *Kocuria rhizophila* | 0.0135 | 98.70% | 0 | *Kocuria* | *Actinomycetales* | Yes |
| 0909A | GCF_000245335 | *Bacillus mojavensis* | 0.0191 | 98.10% | 0 | *Bacillus* | *Bacillales* | Yes |
| 1020B | GCF_029524215 | *Kocuria oceani* | 0.0198 | 98.00% | 0 | *Kocuria* | *Actinomycetales* | No |
| 0102A | GCF_002153395 | *Bacillus subtilis*<br>Group | 0.0208 | 97.90% | 0 | *Bacillus* | *Bacillales* | No |
| Iso2_2024 | GCF_001580365 | *Kocuria turfanensis* | 0.0211 | 97.90% | 0 | *Kocuria* | *Actinomycetales* | Yes |
| 0516A | GCF_007858255 | *Pseudomonas E rhodesiae* | 0.023 | 97.70% | 0 | *Pseudomonas_E* | *Pseudomonadales* | Yes |

**Table 2.** Top three most similar species to 0102A and 0209A isolates identified by autoMLST2. Each organism’s top three species matches are listed in the “Reference Name” column and the sequence ID’s are listed under “Reference Assembly ID”. For each match, the genus, order, and whether or not the match was a type strain, are listed in columns at the right. The quality of the match is assessed by Mash distance (a similarity assessment utilizing mutation distance and P-value significance), average estimated nucleotide identity (ANI-percent nucleotide similarity between two sequences), and P-value (statistical chance that the match was made by chance). 0102A matches are listed in the gray rows and 0209A matches are listed in the white rows.

| Query<br>Organism | Reference<br>Assembly ID | Reference<br>Name | Mash<br>Distance | Estimated Average<br>Nucleotide Identity<br>(ANI) | P-value | Genus | Order | Type<br>Strain |
| --- | --- | --- | --- | --- | --- | --- | --- | --- |
| 0102A | GCF_000332645 | *Bacillus inaquosorum* | 0.0596 | 94.00% | 0 | *Bacillus* | *Bacillales* | Yes |
| 0102A | GCF_004116955 | *Bacillus vallismortis* | 0.0622 | 93.80% | 0 | *Bacillus* | *Bacillales* | Yes |
| 0102A | GCA_031316495 | *Bacillus spizizenii* | 0.0633 | 93.70% | 0 | *Bacillus* | *Bacillales* | Yes |
| 0209A | GCF_000332645 | *Bacillus inaquosorum* | 0.0598 | 94.00% | 0 | *Bacillus* | *Bacillales* | Yes |
| 0209A | GCF_004116955 | *Bacillus vallismortis* | 0.0613 | 93.90% | 0 | *Bacillus* | *Bacillales* | Yes |
| 0209A | GCA_031316495 | *Bacillus spizizenii* | 0.0656 | 93.40% | 0 | *Bacillus* | *Bacillales* | Yes |

**Figure 6.**
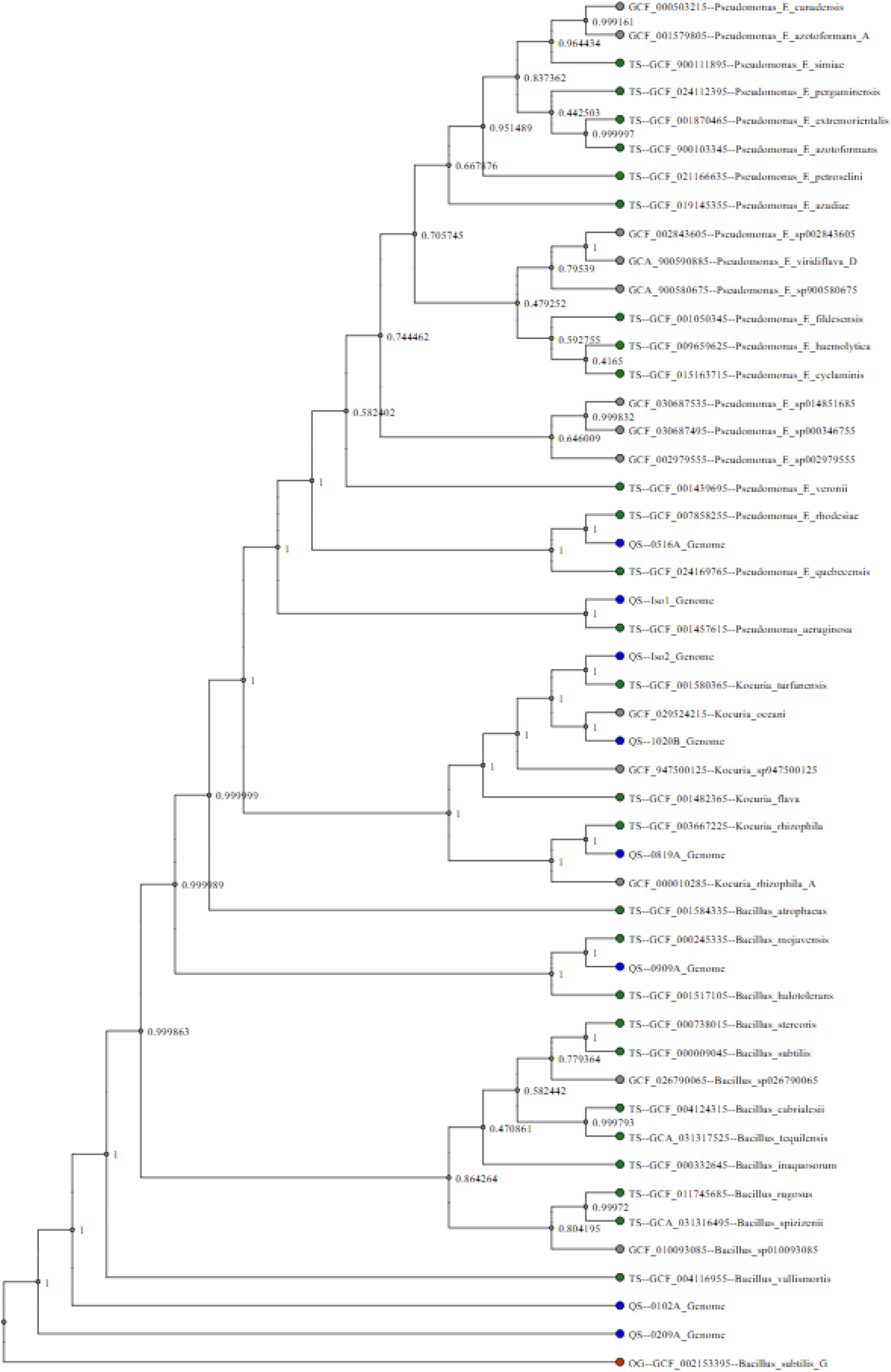
Phylogenetic Tree Generated by autoMLST. This multi-locus species tree was created using all sequenced genomes. autoMLST automatically identified related sequences and aligned them to create a species tree de novo. Query sequences, those from this study, are denoted with blue circles at the ends of their branches. The assigned outgroup is denoted by a red dot on the bottom branch of the tree. All other strains were auto populated when building the tree; type strains are denoted with green dots and all other strains are denoted with gray dots.

All genome FASTA files were analyzed by antiSMASH v8.0.2. (30). antiSMASH identified 5-20 biosynthetic gene clusters (BGC’s) in each genome (Table 3). Potential pigment-producing BGC’s were identified in each genome that was sequenced (Table 4). Pulcherrimin clusters were identified in all *Bacillus sp*. genomes that were sequenced, 0102A, 0209A, and 0909A. Carotenoid clusters were identified in the three *Kocuria sp*. that were sequenced, 0102B, 0819A, and Iso2_2024. Pyoverdine SXM-1 and aryl polyene clusters were identified in 0516A, both of which may contribute to its green color. Pf-5 pyoverdine and pyocyanine clusters were both identified in Iso1_2024. These produce yellow-green and blue pigments, respectively.

**Table 3.** Total number of BGS identified in each genome by antiSMASH.

| Isolate | # of BGC's Identified |
| --- | --- |
| 0516A | 13 |
| 1020B | 7 |
| 0819A | 5 |
| 0102A | 17 |
| 0209A | 20 |
| 0909A | 17 |
| Iso1_2024 | 16 |
| Iso2_2024 | 7 |

**Table 4.** antiSMASH Pigment Cluster Results. antiSMASH identified potential pigment-producing BGC’s in each genome that was sequenced. Pulcherrimin clusters were identified in 0102A, 0209A, and 0909A. Carotenoid clusters were identified in 0102B, 0819A, and Iso2_2024. Pyoverdine SXM-1 and aryl polyene clusters were both identified in 0516A. Pf-5 pyoverdine and pyocyanine clusters were both identified in Iso1_2024.

| Strain | Contig | From | To | Similarity Confidence | Cluster Type | Color |
| --- | --- | --- | --- | --- | --- | --- |
| 0102A | 3 | 317,989 | 338,834 | High | pulcherriminic acid | brown-red |
| 0209A | 2 | 625,311 | 646,057 | High | pulcherriminic acid | brown-red |
| 0909A | 3 | 260,785 | 281,531 | High | pulcherriminic acid | brown-red |
| 1020B | 1 | 3,314,706 | 3,356,792 | Low | carotenoid | yellow |
| 0819A | 1 | 761,663 | 786,188 | Low | carotenoid | yellow |
| Iso2_2024 | 1 | 1,202,265 | 1,246,433 | Low | carotenoid | orange |
| 0516A | 1 | 1,660,402 | 1,749,408 | Low | pyoverdine SXM-1 | yellow-green |
| 0516A | 1 | 5,956,127 | 5,999,702 | Low | aryl polyene | yellow |
| Iso1_2024 | 1 | 3,827,676 | 3,961,195 | Low | Pf-5 pyoverdine | yellow-green |
| Iso1_2024 | 1 | 4,502,360 | 4,523,372 | High | pyocyanine | blue |

Biosynthetic gene cluster analysis by antiSMASH identified that each *Kocuria* strain that was sequenced produced carotenoid pigments (Table 4) (30). To dig deeper into what types of carotenoids each strain produces, DIAMOND software was used to identify individual carotenoid genes (31). It was determined that strains 0819A and 1020B produce yellow pigments that are like decaprenoxanthin and that strain Iso2_2024 produces an orange carotenoid closely related to bacterioruberin (Figure 7). Predicted colors of intermediates and products are depicted in small squares to the left of each pathway in Figure 7 that are colored either cream, yellow, orange, or red. Enzyme confidence is indicated by placing enzyme names in either green rectangles (high confidence) or orange rectangles (medium confidence). Confidence level was determined based on a combination of qualitative and quantitative assessments, factoring in percent identity matches, annotation name of the aligned protein, genome positioning relative to other genes in the pathway, and supporting literature from the *Kocuria* genus and species-specific carotenoid production.

**Figure 7.**
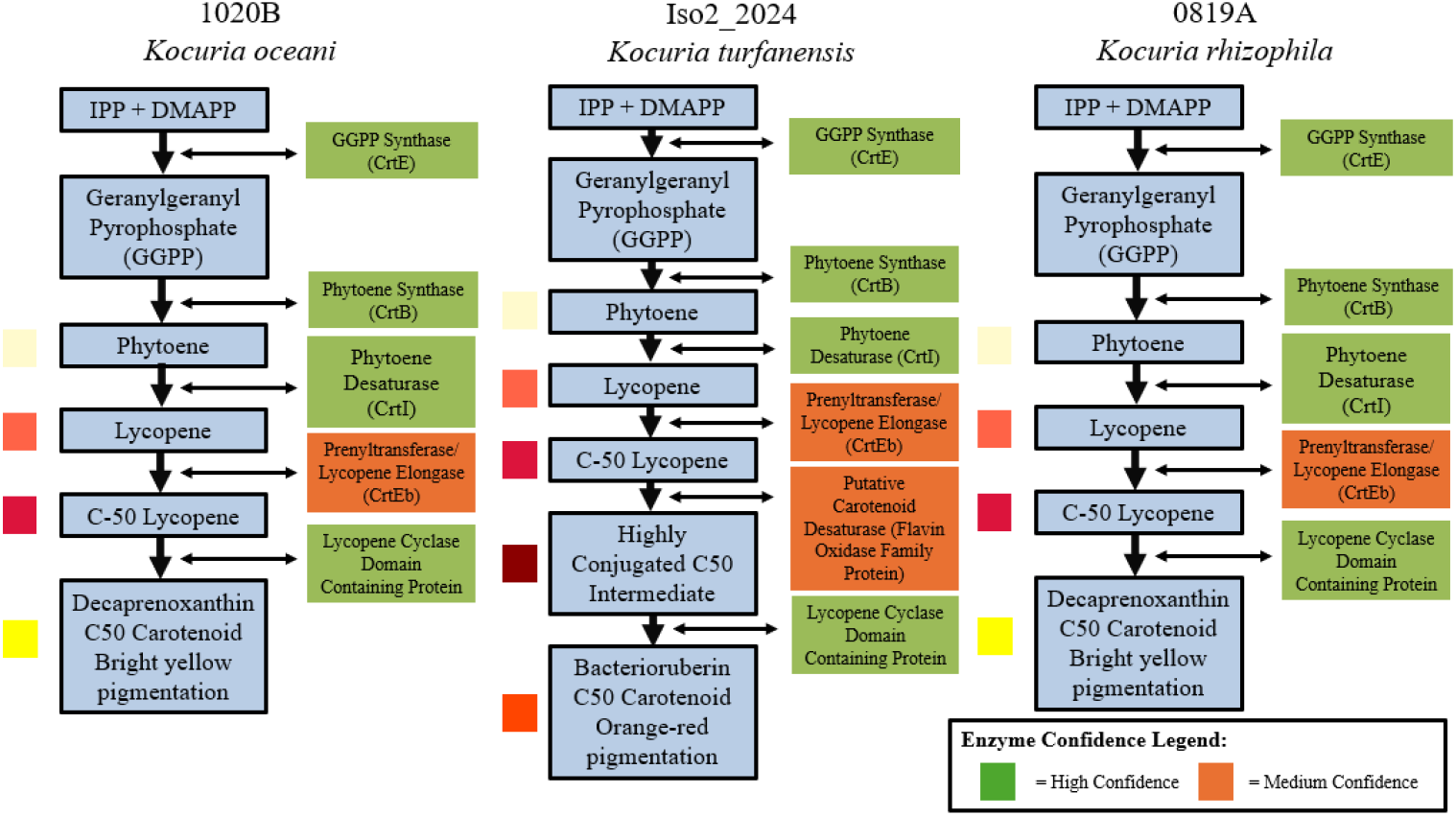
Theoretical pathways of carotenoid-producing Biosynthetic Gene Clusters in three *Kocuria* strains. *K. oceani* strain 1020B produces a yellow pigment like decaprenoxanthin. (left) *K. turfanensis* produces an orange pigment like bacterioruberin. (middle) *K. rhizophila* strain 0819A produces a yellow pigment like decaprenoxanthin. (right) The intermediates and products that are produced through each pathway are shown in blue rectangles. The enzymes that perform each reaction are outlined in either green or orange, depending on the confidence of the prediction. The colors of the intermediates and products are shown in colored squares, either cream, orange, red, or yellow, to the left of their respective blue rectangles.

antiSMASH also identified antibiotic and/or bioactive compound BGC’s in each genome that was sequenced (Table 5) (30). The *Bacillus sp*. were found to produce a wide range of antibiotic compounds including fengycin, bacillaene, macrolactin H, bacilysin, subtilosin A, bacillibactin, zwittermicin A, plipastatin, staphylococcin C55, and surfactins. The *Kocuria sp*. were each found to produce siderophores. The identified *Pseudomonas sp*. produce different varieties of bioactive compounds. 0516A, *P. oceani*, was shown to produce pyoverdine SXM-1, pyochelin, a betalactone, and ambactin. Iso1_2024, *P. aeruginosa*, was shown to produce pseudopaline, pyochelin, azetidomonamide, Pf-5 pyoverdine, L-2-amino-4-methoxy-trans-3-butenoic acid, hydrogen cyanide, pyocyanine, and a homoserine lactone.

**Table 5.** Identification of bioactive compound BGCs by antiSMASH in each genome sequence. All genomes are predicted to produce possible bioactive compounds. Results from each strain are clustered into rows and separated using gray and white colors for ease of visibility.

| Strain | Contig | From | To | Similarity | Cluster Type |
| --- | --- | --- | --- | --- | --- |
| 0102A | 2 | 1 | 27,421 | Medium | fengycin |
| 0102A | 2 | 94,777 | 209,569 | High | bacillaene |
| 0102A | 2 | 473,171 | 563,633 | High | macrolactin H |
| 0102A | 3 | 31,390 | 72,808 | High | bacilysin |
| 0102A | 3 | 78,239 | 99,852 | High | subtilosin A |
| 0102A | 5 | 167,049 | 233,315 | High | bacillibactin |
| 0102A | 6 | 183,218 | 264,567 | Low | zwittermicin A |
| 0102A | 8 | 177,022 | 242,413 | High | surfactin |
| 0102A | 16 | 1 | 12,827 | Low | plipastatin |
| 0209A | 1 | 1 | 80,039 | High | fengycin |
| 0209A | 1 | 129,022 | 243,837 | High | bacillaene |
| 0209A | 1 | 475,315 | 565,767 | High | macrolactin H |
| 0209A | 2 | 88,066 | 115,943 | High | staphylococcin C55 $\alpha$/staphylococcin C55 $\beta$ |
| 0209A | 2 | 335,497 | 376,915 | High | bacilysin |
| 0209A | 2 | 379,906 | 401,519 | High | subtilosin A |
| 0209A | 2 | 920,550 | 986,881 | High | bacillibactin |
| 0209A | 3 | 1 | 22,199 | Low | plipastatin |
| 0209A | 4 | 212,369 | 293,721 | Low | zwittermicin A |
| 0209A | 5 | 194,941 | 220,118 | Low | surfactin |
| 0209A | 7 | 1 | 27,593 | Low | surfactin |
| 0209A | 12 | 1 | 13,658 | Low | plipastatin |
| 0209A | 13 | 1 | 10,500 | Low | plipastatin |
| 0909A | 2 | 1 | 27,607 | High | fengycin |
| 0909A | 3 | 528,659 | 550,271 | High | subtilosin A |
| 0909A | 3 | 554,481 | 595,899 | High | bacilysin |
| 0909A | 5 | 196,546 | 222,617 | Low | surfactin |
| 0909A | 6 | 1 | 27,764 | Low | surfactin |
| 0909A | 7 | 23,680 | 75,454 | High | bacillibactin |
| 0909A | 11 | 1 | 12,886 | Low | plipastatin |
| 1020B | 1 | 3,435,405 | 3,465,789 | Low | siderophore |
| 0819A | 1 | 2,248,980 | 2,278,905 | na | siderophore |
| Iso2_2024 | 1 | 1,344,235 | 1,374,619 | Low | siderophore |
| 0516A | 1 | 1,660,402 | 1,749,408 | Low | pyoverdine SXM-1 |
| 0516A | 1 | 2,115,018 | 2,168,603 | High | pyochelin |
| 0516A | 1 | 3,080,942 | 3,109,289 | na | betalactone |
| 0516A | 1 | 5,598,641 | 5,642,018 | Low | ambactin |
| Iso1_2024 | 1 | 150,340 | 172,429 | High | pseudopaline |
| Iso1_2024 | 1 | 1,788,812 | 1,850,153 | High | pyochelin |
| Iso1_2024 | 1 | 2,788,139 | 2,835,197 | High | azetidomonamide A/azetidomonamide B |
| Iso1_2024 | 1 | 3,827,676 | 3,961,195 | Low | Pf-5 pyoverdine |
| Iso1_2024 | 1 | 4,015,469 | 4,067,749 | High | L-2-amino-4-methoxy-trans-3-butenoic acid |
| Iso1_2024 | 1 | 4,165,722 | 4,178,683 | High | hydrogen cyanide |
| Iso1_2024 | 1 | 4,502,360 | 4,523,372 | High | pyocyanine |
| Iso1_2024 | 1 | 5,016,612 | 5,037,217 | na | homoserine lactone |

Each genome from this study, along with similar strains from other environments, was analyzed by the G4 Hunter online server (32,47,48). The *Kocuria* strains from this study had the highest G-quadruplex (G4) frequencies, above 4 occurrences per thousand base pairs (kb), or 11,606-19,373 total G4’s. (Table 6) This was slightly higher than *K. rhizophila* strain 7_17 (GenBank ID: CP124833.1) which contained 10,368 G4’s at a frequency of 3.8 per 1kb. The *Pseudomonas sp*. have comparatively lower G4 content. Strains 0516A and Iso1_2024 have a G4 frequency of 1.5 per 1kB with a total of 9,072 or 9,632 total G4’s, respectively. These were slightly lower than G4 concentration in *P. aeruginosa* clinical strain AR_0230, which contained 10,336 G4s at a frequency of 1.5 per 1kb (47). The *Bacillus* strains have the lowest estimated G4 content and are each estimated to have a 0.4 per 1kb frequency, or 1,462 – 1,693 G4’s. This is consistent with estimated values for *B. subtilis* strain 3610 (48). These differences in frequency align with the GC content of each genome. The *Kocuria* species had GC contents over 70%, the *Pseudomonas* strains had GC percents in the 60%’s, and the *Bacillus* strains all had GC percents in the 40%’s.

**Table 6.**
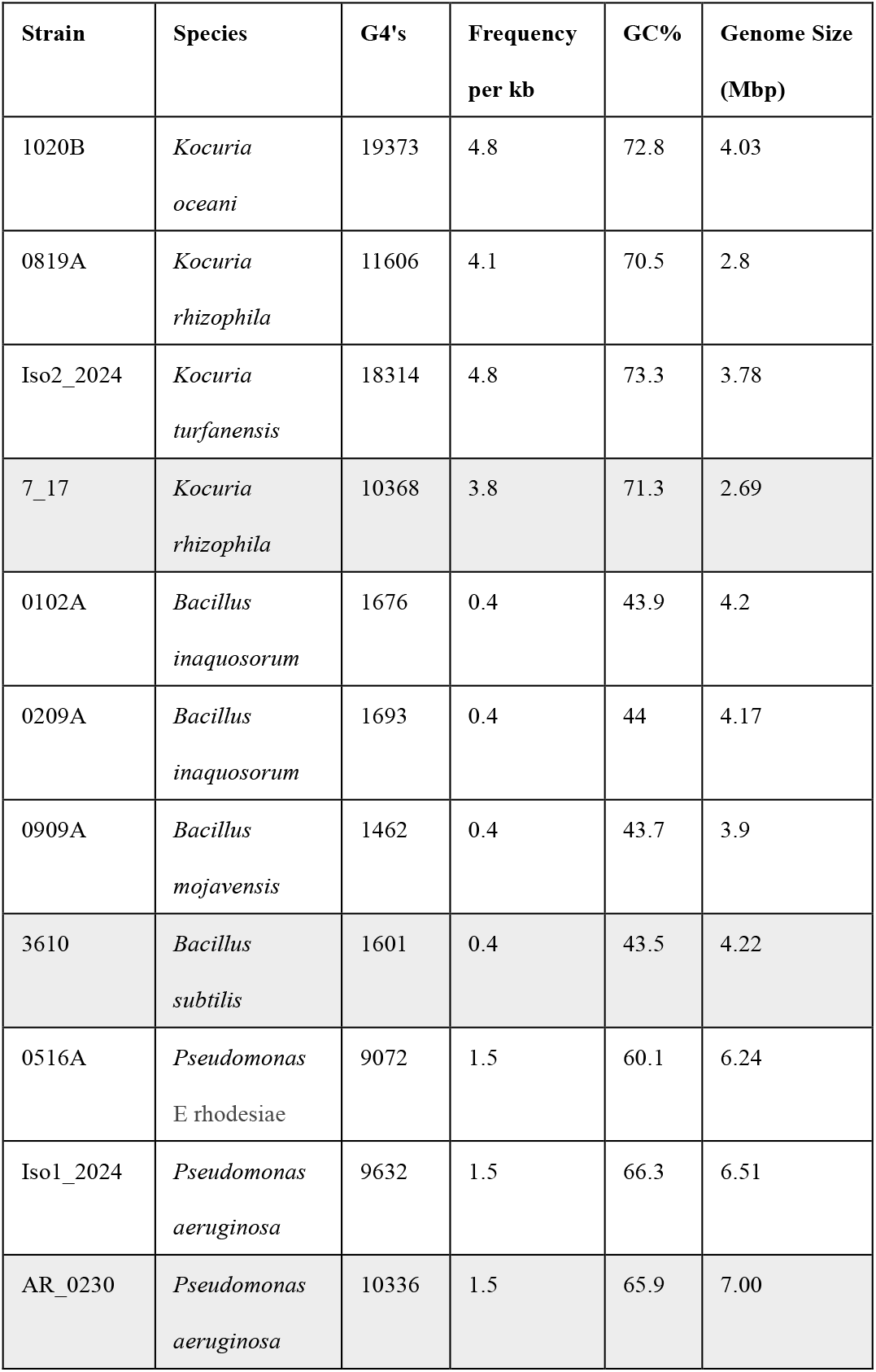
G4s and GC Content of isolates as assessed by G4 Hunter. Sequence files of environmental strains from this study (white rows) and similar strains from other environments (gray rows) were assessed by the G4 Hunter webserver.

## 4. Discussion

In this study, we used sequencing and bioinformatics to assess microbial diversity and identify mechanisms that are beneficial for bacterial survival in inhospitable environments. Using the wide-range method of Illumina 16S rRNA sequencing, we identified the dominant phyla to be *Actinobacteria* and *Proteobacteria* in Atacama soil (Figure 3B). As expected, the locations with more plant life had a higher abundance of bacteria, and the two salt flat areas, Chaxa and Tebenquiche, were within the two outlier groups from the rest of the samples (Figure 3A and 3C).

Eight isolates were selected that produce pigments to whole-genome sequence (Figure 5). Using autoMLST2, three *Kocuria* strains were identified which are closely related to *K. rhizophila, K. turfanensis*, and *K. oceani* (Table 1) (29). Two *Pseudomonas* strains were identified which are closely related to P. *aeruginosa* and *P. E rhodesiae*. Three *Bacillus* strains were identified, two of which are closely related to *B. inaquosorum* and one of which is related to *B. mojavensis* (Table 2). Previously, both *B. inaquosorum* isolates were identified as *B. vallismortis* using autoMLST v1 (49). *B. inaquosorum* was previously considered a subspecies of *B. subtilis* (50), which is very closely related to *B. vallismortis* (51). autoMLST2 improves upon autoMLST1 by including new databases of sequence data to create alignments. This added information slightly altered the identification of these bacteria.

Potential pathways for producing the pigments that were seen on a plate were identified bioinformatically (Figure 5 and Table 4). The types of pigments identified have different roles in species survival. Carotenoids often exhibit strong antioxidant activity and confer UV resistance (5). They are a class of tetraterpenoids made through the assembly of 8 isoprene units to make a 40 carbon (C_40_) backbone. A major differentiating characteristic is their conjugated double bond system which absorbs light between 400 and 550 nm in the UV-Vis range. Aryl polyene has a very similar structure and functionality to carotenoids (52). They serve as antioxidants and have a yellow/green color. Pulcherrimin is a red-brown pigment produced by *B. subtilis* and other bacteria and yeast that controls biofilm aging and disassembly (45). It has iron chelating activity and protects the cells from oxidative stress by lowering the levels of reactive oxidative species (ROS). Pyoverdine and pyocyanine are blue/green pigments that are secreted into the environment. These pigments are siderophores, secreted to chelate mainly iron and other metal ions from the environment for cellular processes (53). Pyocyanin also acts as an antimicrobial (54,55). These pigments also act as signaling molecules for virulence factors and are quorum sensors (56). In the low nutrient R2 media there is abundant pigment production, visible within 24 hours after plating, since metals are in low abundance in minimal medium. Pulcherrimin and pyoverdine pigments also play roles in biofilm formation in *Bacillus* and *Pseudomonas sp*., respectively (45,57). Azetidomonamide has also been shown to control biofilm formation in *Pseudomonas* (58). Biofilm has been shown to play a role in survival of bacteria by providing a hydrated coating (matrix) and providing a physical barrier against UV (6).

Potentially bioactive compounds were identified in every species that was sequenced (Table 5). Production of antibacterial compounds allows for bacterial communities to ward off other species and protect their limited resources in a low-nutrient environment. Bioactive compounds were especially abundant in the *Bacilli sp*. (Table 5). This is not unexpected, since several antibiotics were discovered from *B. subtilis* and related species (59–61) Fengycin, bacillaene, bacilysin, subtilosin A, bacillibactin, plipastatin, and surfactin, which were identified in the *B. mojavensis* and B. *inaquosorum* strains, are known to be produced by *B. subtilis* group organisms (Table 5) (60–65). It was predicted that both *B. inaquosorum* strains may produce zwittermicin A, which is unexpected since it is known to be produced by *Bacillus cereus*, which is not part of the *B. subtilis* group. It is also unexpected that strain 0209A, *B. inaquosorum*, is predicted to produce staphylococcin C55, known to be produced by *Staphylococcus aureus* strain C55 (66). Each *Kocuria* strain was shown to produce siderophores, essential for iron uptake, which hold potential for antimicrobial activity (67). *P. oceani* and *P. aeruginosa* strains were also shown to produce several siderophores, including pyoverdine, pyochelin, and polycinine (53). Iso1_2024, *P. aeruginosa*, is capable of producing pseudopaline, an iron chelator for zinc uptake (53). Iso1_2024 is also capable of producing two additional bioactive compounds: l--2-amino-4-methoxy-trans-3-butenoic acid, an antimicrobial, and hydrogen cyanide, which is a generally toxic compound that helps suppress plant fungal diseases (68,69). 0516A, *P. oceani*, produces two additional antimicrobial compounds: one beta lactone and ambactin (Table 5). Closely related *Pseudomonas* species are known to produce beta lactone antibiotics, such as *P. fluorescens* which produces obafluorin (70). Ambactin is infrequently described in literature, but has been identified in other genera, including *Streptomyces* and *Ochrobactrum* (71,72). It has been shown to have anti-MRSA activities in some species. Outside of ambactin, all other compounds produced by 0516A and Iso1_2024 are expected to be produced by *Pseusomonas sp*.

Several strains had very high GC content and high frequency of G4 formation (Table 6). Increased GC content may increase the stability of DNA since guanines and cytosine form more favorable nucleotide stacking patterns than adenine and thiamine (14). Areas that are GC-rich are sometimes capable of forming G4 quadruplexes, a non-helical DNA secondary structure (16). It has been shown that in some bacteria, G4’s regulate RecA-dependent DNA repair pathways (17). This shows that G4 formation could play a role in helping organisms survive in a highly irradiated environment. While the strains from the Atacama didn’t have significantly different GC content or G4 count from similar strains from less extreme environments (Table 6), these features may aid in maintaining DNA stability. Future work will include a more in-depth analysis of G4 placement in these genomes as opposed to the high-level overview provided here.

## 5. Conclusions

Using our genomic sequences, we can identify mechanisms that are beneficial for survival in inhospitable environments. Isolates subjected to WGS were selected for their pigment-production properties, but sequence data allowed us to identify other adaptations including production of antimicrobial and bioactive compounds, key biofilm formation genes, molecules used for nutrient sequestering, and high GC content. The knowledge gained from this survey will enable evolutionary understanding of key survival strategies in bacteria. Insight into how bacteria handle stress is relevant in many fields including agriculture, biotechnology and health.

## Supplementary Materials

The following supporting information can be found at: www.sciepublish.com/xxx/s1

Table S1: Locations, coordinates, and elevations of metagenomic sampling locations.

Table S2: Locations, coordinates, and elevations of sampling locations where bacteria where isolated from for WGS.

## Acknowledgments

We would like to thank all Northeastern University (NU) students who participated in Dialogue of Civilizations (DOC) trips to Chile (2018, 2022, 2024). These students helped perform sample collection and bacterial isolation. Thank you to Joey Lehman Morris for co-teaching these DOC courses and participating in sample collection. Thank you also to our OneSeed team, including Sofia Mardones and Guillermo Maluenda, for guiding us to sampling locations. A special thanks also to the Riquelme Lab for supporting our team while in Chile.

## Author Contributions

Conceptualization: VGC, CR, YC, ARP, NTC; Methodology: VGC, NTC, ARP, EI; Formal Analysis: NTC, ARP, EI, EA, AH, MCF; Investigation: NTC, ARP, EI, EA, AH, NT, MCF, MT; Writing-Original Draft: NTC and ARP; Writing-Review & Editing: NTC, ARP, VGC; Supervision: YC, CR, VGC; Project Administration: VGC, YC; Funding Acquisition: VGC, YC

## Data Availability Statement

WGS projects are filed under the following BioSample numbers on NCBI.

| Strain | BioSample Number |
| --- | --- |
| 0102A | SAMN39706723 |
| 0209A | SAMN39706724 |
| 0909A | SAMN39706725 |
| 0516A | SAMN50557742 |
| 1020B | SAMN50557744 |
| 0819A | SAMN50557743 |
| Iso1_2024 | SAMN50557745 |
| Iso2_2024 | SAMN50557746 |

Additionally, we assessed the G4 and GC content of three publicly available strains, listed here.

| NCBI ID | Species |
| --- | --- |
| CP020102.1 | *Bacillus subtilis* |
| NZ_CP027174.1 | *Pseudomonas aeruginosa* |
| CP124833.1 | *Kocuria rhizophila* |

